# Histone acetylation alters nuclear morphology and architecture of human mesenchymal stem cells in rigidity dependent manner

**DOI:** 10.1101/2025.05.12.652695

**Authors:** Rohit Joshi, Darshan Shah, K.V. Venkatesh, Abhijit Majumder

**Author notes:** **Corresponding Author Abhijit Majumder**- Department of chemical Engineering, Indian institute of technology Bombay, Maharashtra 400076.

## Abstract

Nuclear morphology which is closely linked to various cell functions and disease states, is known to be modulated by mechanical stimuli and chromatin modifications. However, the combined effects of substrate rigidity and chromatin modifications remain largely unexplored. In this study, we investigated the interplay between substrate rigidity and histone acetylation, mediated by the histone deacetylase inhibitor (HDACi) valproic acid (VA), in modulating nuclear morphology in human mesenchymal stem cells (hMSCs) cultured on substrates with varying rigidities. We found that VA-induced chromatin decondensation diminished the effects of soft substrates on nuclear morphological parameters, a change not observed on stiff substrates. Furthermore, we showed that the reduction in nuclear stiffness with VA treatment on both substrates was independent of lamin expression, highlighting chromatin decondensation as a key determinant of the observed change in nuclear stiffness. In addition, we demonstrated that soft substrates induced significant nuclear wrinkling, which was alleviated by HDACi treatment, while no such effect of VA was observed on stiff substrates. Together, our findings suggest that chromatin remodeling via HDACi can override substrate-dependent nuclear mechanotransduction on soft substrates, offering new insights into the epigenetic regulation of nuclear mechanics with potential implications for pathologies associated with nuclear morphology.

## Introduction

Nuclear morphology and mechanics are dynamic indicators of the cellular state, function, and health. Morphometric parameters such as nuclear shape, size, chromatin organization, and rigidity vary significantly across physiological and pathological conditions, including aging, cancer, and stem cell differentiation^1–4^. For example, cancer cells frequently exhibit irregularly shaped nuclei that are correlated with genomic instability, whereas nuclear size increases with cellular aging and senescence^5–9^. Changes in nuclear rigidity also critically influence cellular invasion through complex three-dimensional extracellular matrices, such as during metastasis^10–12^. Additionally, nuclear rigidity increases during stem cell differentiation^4,13^. These observations established the morphometric parameters of the nucleus as biomarkers of cellular state, function, and health.

Nuclear morphometric parameters are influenced not only by biochemical cues, but also by mechanical cues. Among various mechanical cues, substrate rigidity is a well-established regulator of nuclear morphology and chromatin organization^14,15^. Cells cultured on soft substrates exhibit spherical nuclei with condensed chromatin and reduced projected area and volume, whereas those cultured on stiff substrates display flattened nuclei with a more open chromatin structure^14,15^. Nuclear morphology is also strongly influenced by chromatin modifications, such as histone acetylation, which reduces chromatin packing or condensation, thereby increasing gene accessibility^16,17^. Interestingly, chromatin condensation and the resulting epigenetic modifications can also be achieved by culturing cells on substrates with varying stiffness, suggesting an intrinsic link between the mechanical and epigenetic regulation of nuclear structure.

Although pairwise interactions between substrate rigidity, histone acetylation, and nuclear morphology have been explored, the collective interaction of these three parameters remains poorly understood. Here, we investigated a fundamental question: how does chromatin decondensation induced by histone acetylation modifiers influence nuclear morphology and mechanics in microenvironments with varying stiffness? Specifically, we investigated whether chromatin unpacking alters nuclear shape, size, and rigidity differently on soft versus rigid substrates.

To address this question, we cultured human mesenchymal stem cells (hMSCs) on polyacrylamide (PAA) gels of two physiologically relevant stiffnesses (3 and 34 kPa) and treated them with sub-IC50 concentrations of the histone deacetylase inhibitor (HDACi) valproic acid (VA). We found that VA-induced histone acetylation occurred on both soft and stiff substrates, but its effects on nuclear morphology, such as changes in shape, size, and wrinkling, were observed only on soft substrates. Furthermore, VA treatment decreased nuclear rigidity on both substrates without influencing nuclear lamina protein expression, a known key regulator of nuclear rigidity, indicating that the nuclear softening observed upon VA treatment results solely from chromatin opening.

Altogether, our findings reveal that chromatin remodeling via HDACi can override the mechanical regulation of nuclear morphology in soft tissue-like microenvironments. Given that VA is an FDA-approved drug for epilepsy, bipolar disorder, and cancer^18^, these results provide insights into the mechanobiology of chromatin organization and its potential implications in disease pathology and drug response.

## Material and methods

### Substrate preparation

Polyacrylamide gels with elastic moduli of ∼3 and ∼34 kPa were prepared by crosslinking 40% polyacrylamideand 2% bis-acrylamide solutions. The protocol for substrate preparation and the elastic modulus value were adopted from a previously reported study ^19^. Briefly, the gel solution for desired stiffness (∼3 kPa and ∼34 kPa, Table S1) was mixed with ammonium persulfate (1:100) and TEMED (1:1000), and a drop of 130 µL was placed between two glass coverslips, one coated with 3-APTMS (Sigma) and the other with a hydrophobic coating. After polymerization, the hydrophobic coverslips were removed. The gel was coated with type I collagen (25 µg/ml) (Invitrogen; A1048301) using sulfo-SANPAH-based conjugation and kept at 4°C overnight.

### Cell culture

Bone marrow-derived hMSCs were purchased from Lonza (Cat. No. #PT-2501). hMSCs were cultured in low-glucose DMEM (Himedia; AL006) supplemented with 16% FBS (Himedia; RM9955), 1% Antibacterial-Antimycotic (Himedia; A002), and 1% Glutamax (Gibco; 35050) under humidified conditions at 37°C with 5% CO2. The cells were trypsinized with TrypLE™(Gibco; 12604021) once they reached 70% confluency. The cells were seeded on polyacrylamide gels at 2000 cells/cm^2^ in 50 µL of media and flooded after 45 min.

### Treatment with HDACi and inhibitors

To check the effect of HDACi on soft substrate, 0.5mM valproic acid (PHR 1061) was added to the growth media at the time of media flooding. To investigate the effect of cellular contractility on nuclear wrinkling, lysophosphatidic acid (30µM, LPA, Sigma, Cat. No. L7260), and Latrunculin B (1µM, LatB, Calbiochem, Cat. No. 428020) were added to the growth media 24 h after cell seeding for 20 min.

### Immunofluorescence staining

Cells were fixed with 4% paraformaldehyde (PFA) in PBS for 15 min at room temperature (RT) and washed thrice with PBS. Cells were then permeabilized with permeabilizing buffer (0.5% Triton X-100-Sigma Aldrich in CSB) for 10 min and blocked with bovine serum albumin (BSA) (4% bovine serum albumin in PBS) for 30 min to minimize nonspecific protein binding. Anti-lamin A (1:500; mouse; Abcam, Cat. no. 8980), anti-lamin B (1:500, rabbit; Abcam, Cat. No. 16048), Anti-Ack (1:500, rabbit, Abcam, Cat. No. 190479) primary antibodies in 4% bovine serum albumin (BSA) were added to the samples and incubated overnight at 4°C. The primary antibodies were removed, and the samples were rinsed twice with PBS for 10 min. The samples were then incubated at room temperature with secondary antibodies (1:500, donkey anti-mouse Alexa Fluor 488 Cat. No. 21202), donkey anti-rabbit AlexaFluor 569 Cat. No. ab175470) phalloidin (1:400, AlexaFlour 532, Cat. No. A22282), and Hoechst 33342 (Cat. No. H3570) diluted 1:5000 in 4% BSA for 2 h at room temperature. The samples were then rinsed twice with PBS. All immunostained samples were stored in PBS at 4°C until imaging. All samples were imaged at 63x (oil) magnification using a laser scanning confocal microscope (LSM 780, Carl Zeiss).

### Nuclear morphometric parameter calculation

Cells on the gels were imaged using a confocal microscope (LSM, Carl Zeiss) at a 63x oil immersion objective. To visualize the vertical cross-section of the nucleus, z-stacks were obtained, and 3D projections were created using Fiji. Nuclear height was calculated from fluorescence profiles measured at a point on the line drawn along the axis of symmetry of the nucleus cross-section. Z-stacks of confocal fluorescent images were analyzed to calculate the nuclear volume and surface area. Gaussian blur filter was applied to the z-stack to smoothen the staining pattern. Furthermore, a 3D object counter function with a minimal threshold was applied for nuclei separation in Fiji. Nuclei were segmented from Z-stacks using the nuclear edge detector function, and the nuclear volume and surface area were measured^20^.

### Analysis of chromatin condensation parameter (CCP)

DAPI-stained nuclei were imaged using a laser scanning confocal microscope (LSM 780; Carl Zeiss) at 63X (oil). For analysis, individual nuclei were cropped using FIJI to obtain only one nucleus per image. The CCP wascalculated using a MATLAB script, as previously described ^21^, in which a gradient-based Sobel edge detection algorithm was used to measure the edge density of the individual nuclei.

### Coherency analysis

Actin alignment was analyzed using the OrientationJ plugin in ImageJ, which quantifies the coherency of local image features. This plugin was designed to characterize the orientation and isotropic attributes within the region of interest (ROI) by analyzing the structure tensor in the local vicinity. The value of coherency indicates the degree to which the local features are oriented: a coherency value of 1 indicates that the features are uniformly oriented in a single direction, whereas a value of 0 signifies isotropy with no predominant orientation in the analyzed ROI^22^.

### Atomic Force microscopy

The peri-nuclear stiffness of MSCs was determined using atomic force microscopy (AFM). Atomic force measurements were performed using a TR800PB silicon nitride pyramidal tip probe on an MFP-3D (Asylum Research) under contact mode force. The spring constant of the cantilever ranged from 0.09 to 0.27 N m^−1^ and the frequency ranged from 17 to 28 kHz.

### Nuclear Wrinkling Index Calculation

To quantify the visually discernible differences in nuclear morphology on soft versus stiff substrates, an image-processing pipeline was developed for confocal microscopy images of hMSCs nuclei stained with lamin A. The Z-stack maximum intensity projection images from ImageJ were imported into MATLAB to estimate the wrinkling index (WI) parameter from the observed lamin A crowding. The algorithm includes a series of pre-processing steps, such as contrast enhancement, smoothing, and image segmentation, to eliminate image grains and isolate the nuclear region from the rest of the image. This was followed by Sobel edge detection to extract the edges of the nucleus. Finally, the nuclear WI was calculated from the resulting binarized edge map as the ratio of the edge pixels to the total nucleus pixels in the image, expressed as a percentage.

### Statistical analysis

Data are presented as mean ± standard deviation (otherwise mentioned) and were analyzed using Microsoft Excel software. Data were plotted using the OriginLab software (IIT Bombay License). Statisticallysignificant differences are indicated by *p < 0.05.

## Results

### HDACi Valproic Acid (VA) increases histone acetylation in a rigidity dependent manner

While it is obvious that a histone deacetylase inhibitor (HDACi) should hyperacetylate histones, its effects on substrates of different rigidities remain to be explored. To answer this primary question, human mesenchymal stem cells (hMSCs), the epigenetics of which are known to be sensitive to substrate rigidity^23^, were cultured on collagen-coated polyacrylamide (PA) hydrogels of physiologically relevant stiffness (34 kPa as stiff; osteogenic lineage-inducing stiffness and 3 kPa as soft; adipogenic lineage-inducing stiffness). The cells were treated with a sub IC50 concentration of an FDA-approved histone deacetylase inhibitor, valproic acid (VA). After 24 h of treatment, the individual and combined effects of substrate rigidity and HDACi on global histone acetylation were evaluated by measuring the levels of histone acetylation (Ack).

Consistent with earlier findings ^15^, hMSCs cultured on stiff hydrogels showed a significant increase (∼4.8-fold) in global histone acetylation compared to those cultured on soft hydrogels (Fig. 1 A&B (i-iii) and 1C). Furthermore, upon addition of valproic acid, global histone acetylation increased significantly on both soft and stiff substrates. However, the fold change on soft substrates was approximately two times higher (∼2 times) than that on stiff substrates (∼1.2 times) (Fig. 1C). Despite this, the non-inhibitory concentration of VA could not fully reverse the effect of substrate stiffness on histone acetylation i.e. 3kPa+VA still showed significantly less histone acetylation than 34 kPa alone.

**Fig. 1.**
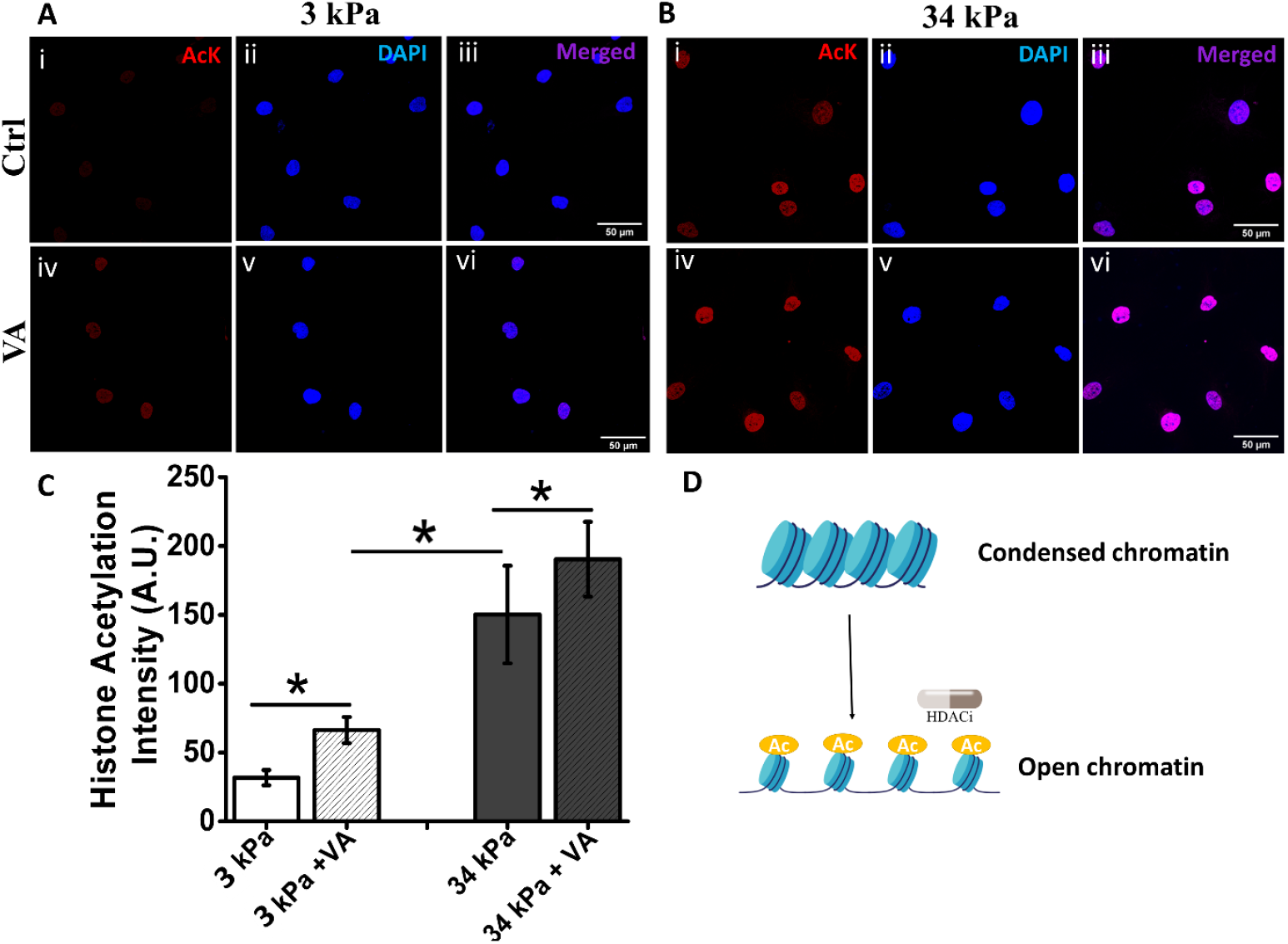
Effect of substrate stiffness and HDACi Valproic Acid (VA) on histone acetylation: VA increases histone acetylation on varying stiffness: Representative immunofluorescence images of hMSCs cultured on (A) soft hydrogel (3 kPa) and (B) stiff hydrogel (34 kPa) without (i-iii) and with (iv-vi) VA. Global Histone acetylation (AcK) (red), nucleus (DAPI) (blue), and merged images (magenta) are shown in (i), (ii), and (iii), respectively. (C) Quantification shows the change in histone acetylation for hMSCs cultured on soft and stiff hydrogels with and without VA. (D) Schematic showing the effect of VA on chromatin compaction. *p<0.05, n>50 nuclei scale bar: 50 µm

### HDACi reduces chromatin condensation in hMSCs cultured on soft substrate to the level of the same on rigid substrate

Next, we asked how the observed histone acetylation resulting from the combined effect of substrate rigidity and HDACi may reflect on chromatin packing. For this purpose, we measured chromatin condensation parameter (CCP), a well-established method for estimating chromatin packing ^15,21,24,25^. Briefly, images of DAPI-stained nuclei were captured at high magnification. The image was converted into a heatmap based on the local intensity of DAPI, in which warmer colors represent condensed or densely packed chromatin (Fig. 2A & B). Calculated CCP values showed ∼1.5-fold higher chromatin compaction on the soft hydrogel as compared to that on the stiff hydrogel (Fig. 2A-i, B-I & C). This typical dependence of chromatin packing on substrate rigidity matched well with the observations of earlier researchers ^26–28^. Furthermore, histone hyperacetylation with VA significantly decreased CCP for both stiff (∼1.3 fold) and soft (∼1.6 fold) substrates (Fig. 2A-ii, B-ii, & C), bringing the CCP on soft with VA down to the level of CCP on a stiff substrate alone.

**Fig. 2.**
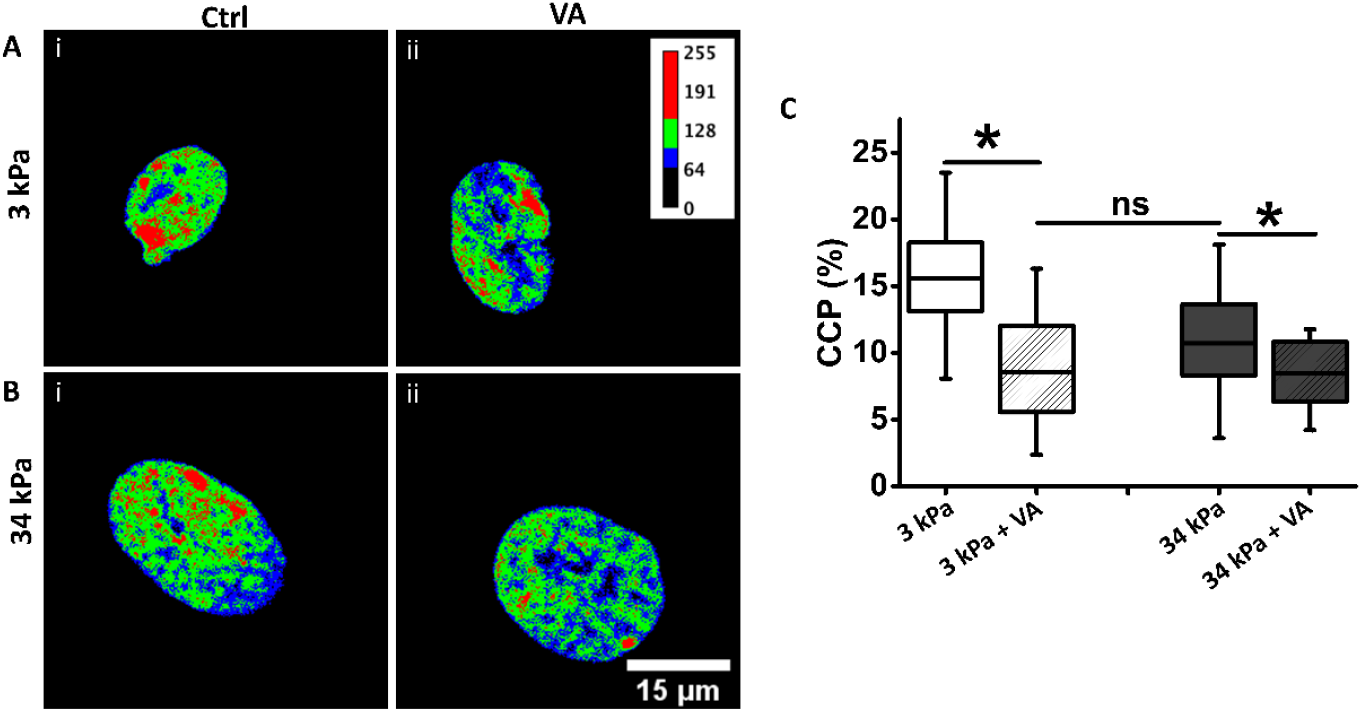
HDACi decreases the chromatin condensation parameter in both stiff and soft substrates: Heat map of DAPI intensity of nucleus on (A) soft hydrogel (3 kPa), (B) stiff hydrogel (34 kPa) without (i) and with (ii) VA, respectively. (C) Quantification of CCP under mentioned conditions. *p<0.05, n>25 nuclei scale bar: 15 µm

### Chromatin decondensation by HDACi changes nuclear morphometric parameters for hMSCs cultured on soft hydrogel but not on stiff hydrogel

In the previous sections, we showed that the effects of VA on global histone acetylation and chromatin condensation are rigidity dependent. Next, we examined how VA-induced chromatin decondensation influenced nuclear morphology on substrates with different stiffnesses. On a soft substrate (3 kPa), valproic acid treatment significantly increased the projected nuclear area by approximately 1.3-fold (Fig. 3A&C), nuclear volume by ∼1.4-fold (Fig. 3D), and surface area by around 1.3-fold (Fig. 3E). At the same time, it reduced the nuclear height by approximately 1.2-fold (Fig. 3F). While reduction in height and increase in projected area can be explained by possible increase in the normal compressive stress acting on the apical surface of the nucleus^29^ due to increased cellular traction upon treatment with VA ^27^, increase in volume indicates an internal outwards stress, probably generated from the opening of the chromatin. Interestingly, on a stiff substrate, no such changes were observed for any of the measured parameters (Fig. 3B, C-F). Further, the nuclear aspect ratio remained unaltered for both rigidity variation and VA treatment, suggesting that the overall elliptical nuclear shape was unaffected by these treatments. (Fig. 3G). Overall, our results suggest that the nuclear morphology of cells present in a soft microenvironment is much more sensitive to chromatin remodeling than that of cells on a rigid substrate.

**Fig. 3.**
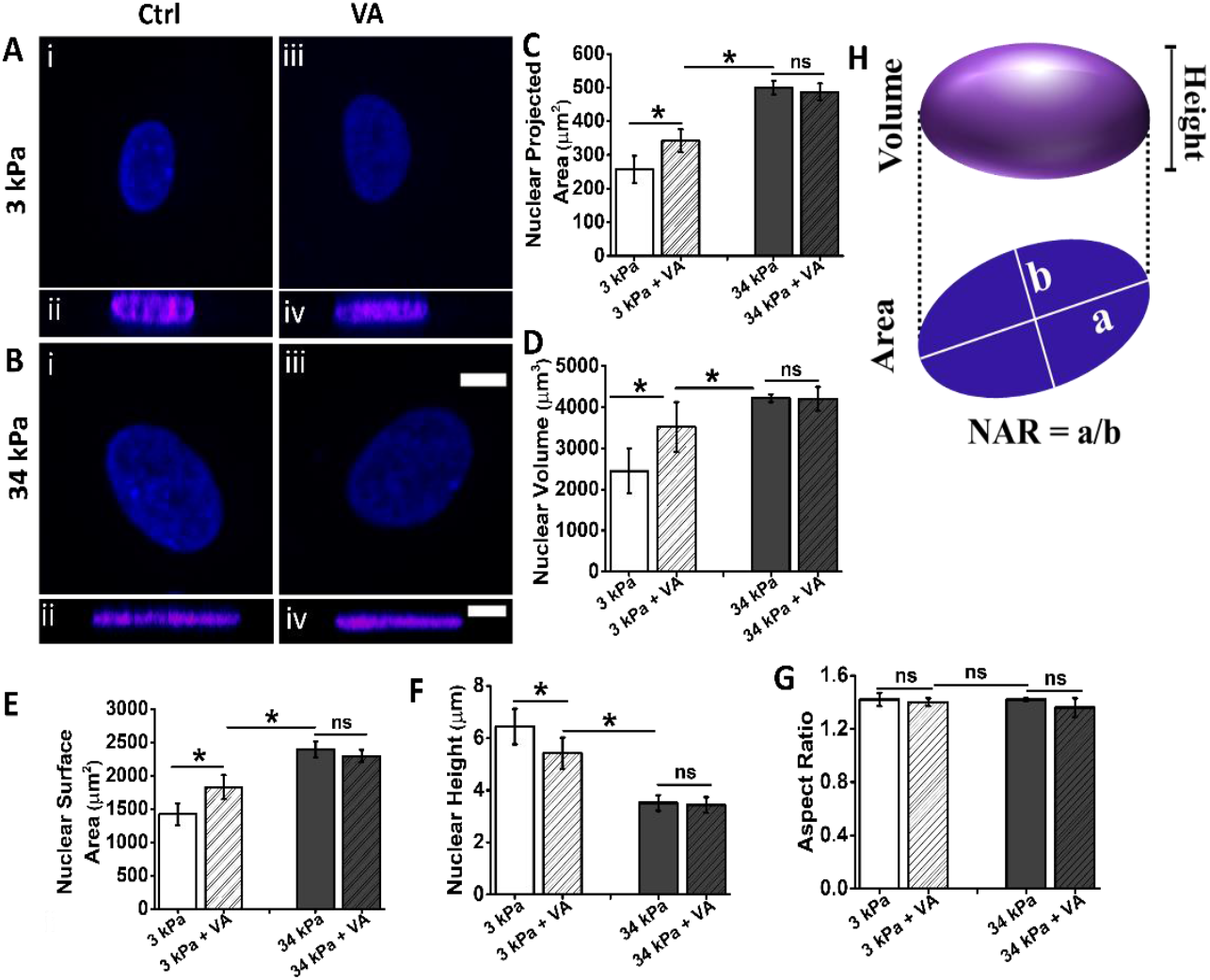
Influence of substrate stiffness and HDACi on nuclear morphology: Representative DAPI stained images of nuclei of hMSCs cultured on (A) soft hydrogel (3 kPa) and (B) stiff hydrogel (34 kPa) without (i) and with (iii) VA, with respective cross-sectional images (ii, iv). Quantification of nuclear morphological parameters (C) nuclear projected area, (D) nuclear volume, (E) nuclear surface area, nuclear height, (G) nuclear aspect ratio, (H) schematic representing quantification *p<0.05, n>50 nuclei scale bar: 10 µm

### Chromatin decondensation decreases nuclear stiffness on soft as well as rigid substrates

Next, we investigated the combined effect of substrate rigidity and VA on nuclear stiffness, which provides mechanical integrity to the nucleus and safeguards it from DNA damage caused by various mechanical stresses^30,31^. Altered nuclear stiffness leads to several pathological conditions, such as progeria, cardiomyopathy, muscular dystrophy, cancer, and senescence ^32,33^. Both substrate stiffness and histone acetylation are known to independently regulate nuclear rigidity^34–36^; however, their combined influence is unknown. To address this question, we examined the combinatorial effect of substrate stiffness and VA on nuclear stiffness using atomic force microscopy (AFM). Measurements were taken after ensuring that the AFM cantilever tip made an indentation just above the nucleus, as shown in the schematic in Fig. 4A and Fig. 4A’. We found that cells cultured on soft substrates showed significantly lower (∼ 1.6-fold) nuclear stiffness than those cultured on stiff substrates (Fig. 4B), which is in agreement with the published literature^34^. Interestingly, unlike other nuclear parameters assessed in this study, which showed no significant response to valproic acid (VA) treatment on stiff substrates, VA consistently reduced nuclear stiffness on both soft and stiff substrates. However, this reduction was more pronounced on the soft substrate, with a decrease of approximately 2.2-fold, compared to a 1.7-fold reduction on the stiff substrate (Fig. 4B).

**Fig. 4.**
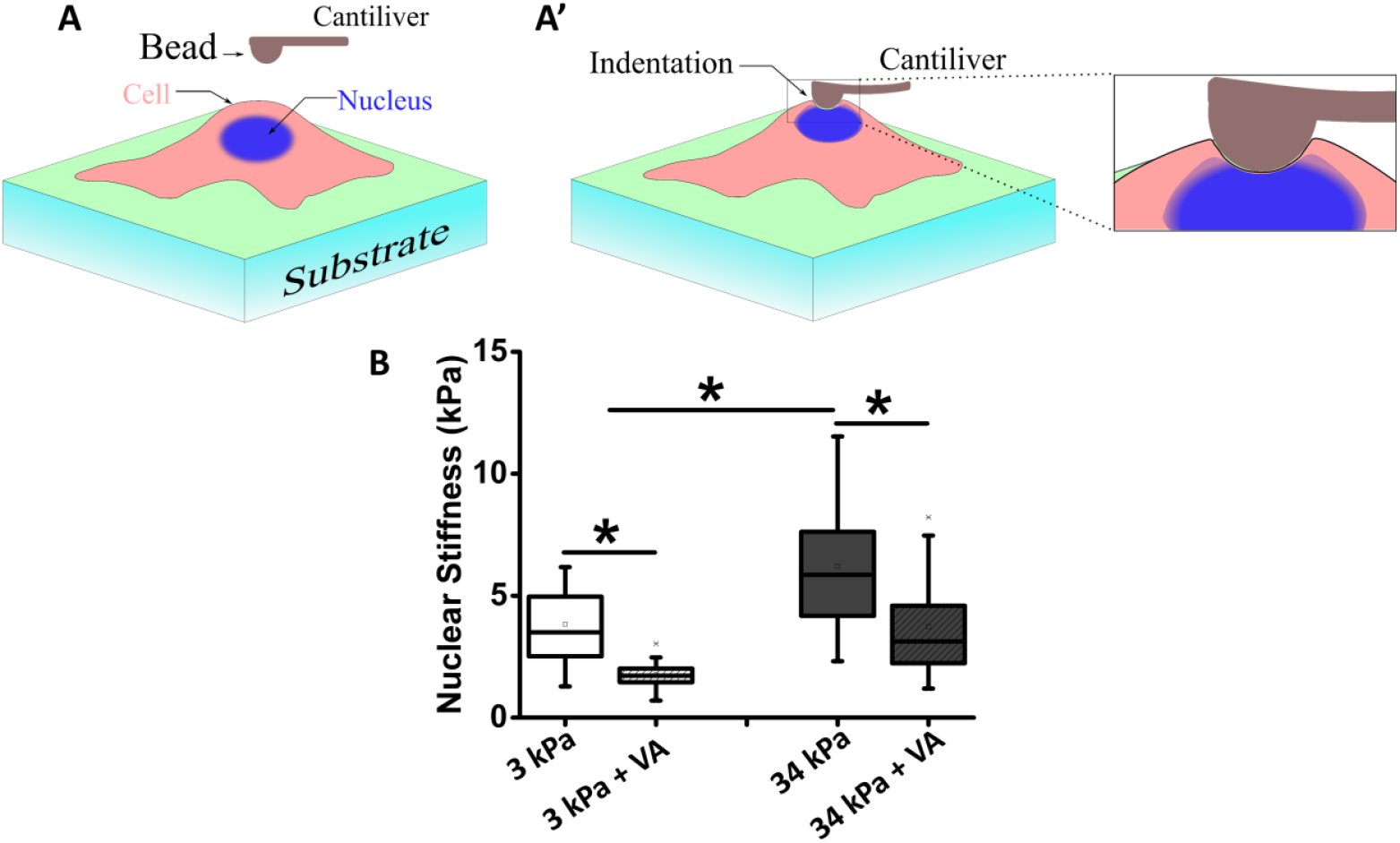
HDACi decreases nuclear stiffness on both stiff and soft substrates: Schematic of (A) AFM setup and (A’) cantilever position on nucleus used for determining nuclear stiffness; (b) Stiffness of nucleus on soft and stiff hydrogels, with and without VA *p<0.05, N=3, n>25 nuclei.

**Fig. 5.**
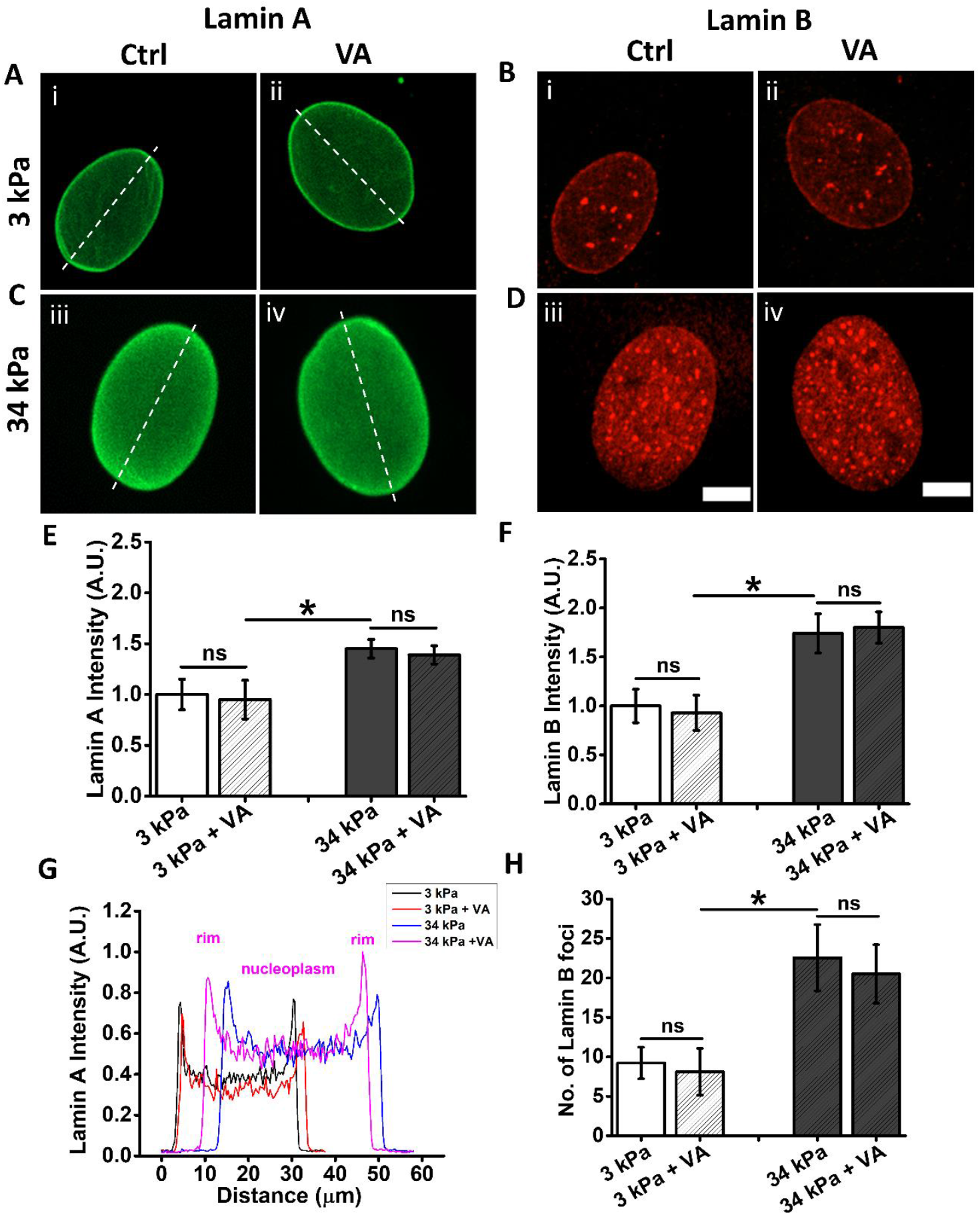
Effect of substrate stiffness and HDACi on Lamin: (A, B) Representative immunofluorescence images of lamin A (A) and lamin B (B) on soft hydrogel (3 kPa) and (C, D) stiff hydrogel (34 kPa) without (i-ii) and with (iii-iv) VA (green – lamin A and Red – lamin B). Quantification of the intensity of Lamin A (E) and lamin B (F) and (G) lamin A distribution in the nucleus along the dashed line. (F) number of foci observed in Lamin B staining. *p<0.05, n>50 nuclei scale bar: 10 µm

### Nuclear lamina proteins lamin A and lamin B are not responsible for HDACi induced decrease in nuclear stiffness

Nuclear stiffness is a combined outcome of chromatin condensation and expression of nuclear lamina proteins lamin A/C and lamin B at the nuclear envelope^37–40^. Increased expression of lamin proteins correlates with greater nuclear stiffness owing to their structural role in resisting deformation. In parallel, highly compact chromatin (heterochromatin) contributes to increased nuclear rigidity, whereas less compact chromatin (euchromatin) reduces nuclear stiffness^39^. In the previous section, we showed that nuclear stiffness increased with an increase in substrate stiffness and decreased upon treatment with VA on soft as well as on rigid substrates. As the expression of lamins is substrate rigidity dependent^41^, to investigate if the observed change in nuclear stiffness upon treatment with VA is associated with a corresponding change in the expression of lamin A and/or lamin B, we estimated the levels of lamin A and lamin B in cells cultured on stiff and soft substrates with and without VA. Immunofluorescence images showed that the cells cultured on the stiff substrate had significantly higher expression of lamin-A (∼1.7 fold) and lamin-B (∼1.5 fold) as compared to cells cultured on the soft substrate (Fig.5 Ai&iii, Bi&iii, E). However, treatment with HDACi did not significantly change the expression of either of the lamins on either of the substrates as compared to control (Fig.5A-F).

As Increased localization of lamin A at the nuclear periphery, combined with enhanced nucleoplasmic expression, is associated with increased nuclear stiffness^37^, we examined the spatial distribution of lamins of varying stiffness with and without VA. On soft substrates, Lamin A showed higher localization along the nuclear periphery with minimal nucleoplasmic distribution, whereas stiff substrates showed higher peripheral as well as nucleoplasmic distributions (Fig.5G). However, VA treatment did not result in any significant changes. Moreover, on soft substrates, lamin B had fewer nucleoplasmic foci than on stiff substrates (Fig.5H). As observed for laminA, here also VA had no influence on either substrate. These observations matched well with the observed changes in nuclear rigidity, indicating that the observed reduction in nuclear stiffness upon VA treatment was due to chromatin decondensation and not due to a change in the expression or distribution of either lamin.

### HDACi decreases nuclear wrinkling on soft hydrogels

Another critical yet less explored nuclear morphometric feature is the wrinkling of the nuclear lamina, which has recently been associated with cancer^42^ and stem cell differentiation^43^. While earlier work has attributed nuclear wrinkling to actin cytoskeletal tension and impingement of the nuclear envelope^43–45^, however how chromatin compaction on substrates of different rigidity influences this feature has not been explored. We observed that hMSCs cultured on soft hydrogel (3kPa) had prominent wrinkles on their nuclear envelope (Fig. 6A), but such wrinkles were rarely observed on the nuclei on stiff gels (Fig. 6B), as observed in the XY, XZ, and YZ projections. For the cells cultured on soft hydrogels, nuclear wrinkling decreased in terms of the percentage of cells with nuclear wrinkles and the wrinkling index (Fig. 6C, E, and F) upon treatment with VA. No such change was observed on rigid substrates (Fig. 6D, E, and F), which correlates with our earlier observation of the increase in nuclear volume and surface area (Fig. 3D & E) upon treatment with VA on soft gels, but not on stiff substrates. Quantitatively, 60% of human mesenchymal stem cells cultured on soft substrates exhibited nuclear wrinkling, which decreased to 40% after VA treatment. In contrast, only 20% of the cells on the stiff substrate displayed weak nuclear wrinkling, and VA treatment had no effect. While the number of cells with wrinkles provides a binary result (yes/no), it does not differentiate between two cells with different degrees of nuclear wrinkling. To quantify this, we used an edge detection algorithm developed by Mauck’s group^43^ and found that cells cultured on soft hydrogels had an overall higher mean wrinkling index (∼1.6-fold) than the nucleus of the cells cultured on stiff hydrogels (Fig. 6F). Upon treatment with VA, cells cultured on soft gels showed a significant decrease in nuclear wrinkling (∼1.2-fold), while no significant change was observed when cultured on stiff gels in the presence of VA. Furthermore, nuclear wrinkling, wherever observed, showed no specific apical/basal polarity, as observed in Z-stack slices (Fig. 6G and H).

**Fig. 6.**
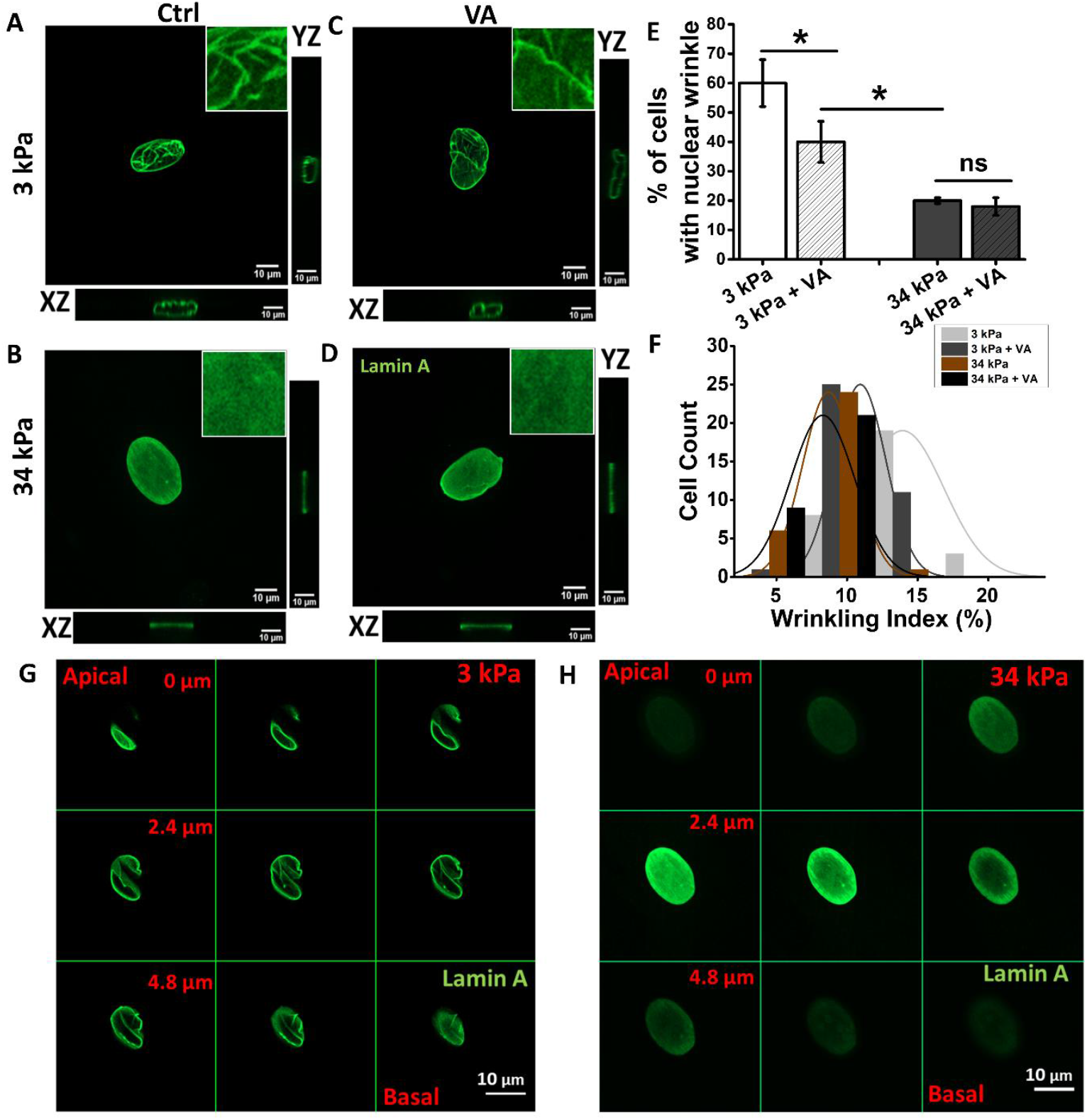
Effect of substrate stiffness and HDACi on nuclear wrinkling: Representative immunofluorescence images showing comparison of nuclear wrinkling in Lamin A-stained nuclei from maximum projection confocal images of hMSCs cultured on stiff (3 kPa) and soft (34 kPa) hydrogels with (A, B) and without VA (C, D). Montage (E) showing corresponding individual z-stack slices of hMSC wrinkling across all planes of the nuclear envelope for cells cultured on soft hydrogel; Graph shows quantification of the percentage of cells with nuclear wrinkles for cells cultured on stiff (3 kPa) and soft (34 kPa) hydrogels with and without VA treatment. Graph (H) shows quantification of wrinkle index for cells cultured on stiff (3 kPa) and soft (34 kPa) hydrogels with and without VA. *p<0.05, n>30 nuclei, scale bar: 10 µm

As hypothesized earlier, nuclear wrinkling results from the availability of excess membrane when the nuclear volume is changed by external or internal factors^44,46^. We further modulated the nuclear volume and surface area by treating hMSCs cultured on soft and stiff hydrogels with the actin depolymerizing agent Latrunculin B (LatB), and on stiff hydrogels with the pharmacological agonist lysophosphatidic acid (known to increase cellular contractility in hMSCs) ^47^. Upon treatment of hMSCs cultured on soft hydrogels with LPA, an increase in nuclear volume and a concomitant (∼1.6-fold) reduction in nuclear wrinkling were observed. In contrast, LatB treatment increased nuclear wrinkling in cells on both stiff and soft substrates (Fig. 7A-C). However, the increase in wrinkling was more prominent for the stiff substrate (∼4-fold) than for the soft one (∼1.5-fold). This is presumably because nuclei on stiff substrates have a larger nuclear surface area and, hence, more available nuclear membranes for folding^44^. Additionally, we found that upon treatment with LPA, hMSCs cultured on soft hydrogels showed (∼1.5-fold higher expression (Fig. S1A) and more coherent actin stress fibers (∼4-fold) (Fig. S1B), two hallmarks of increased cellular contractility, compared to the untreated control.

**Fig. 7.**
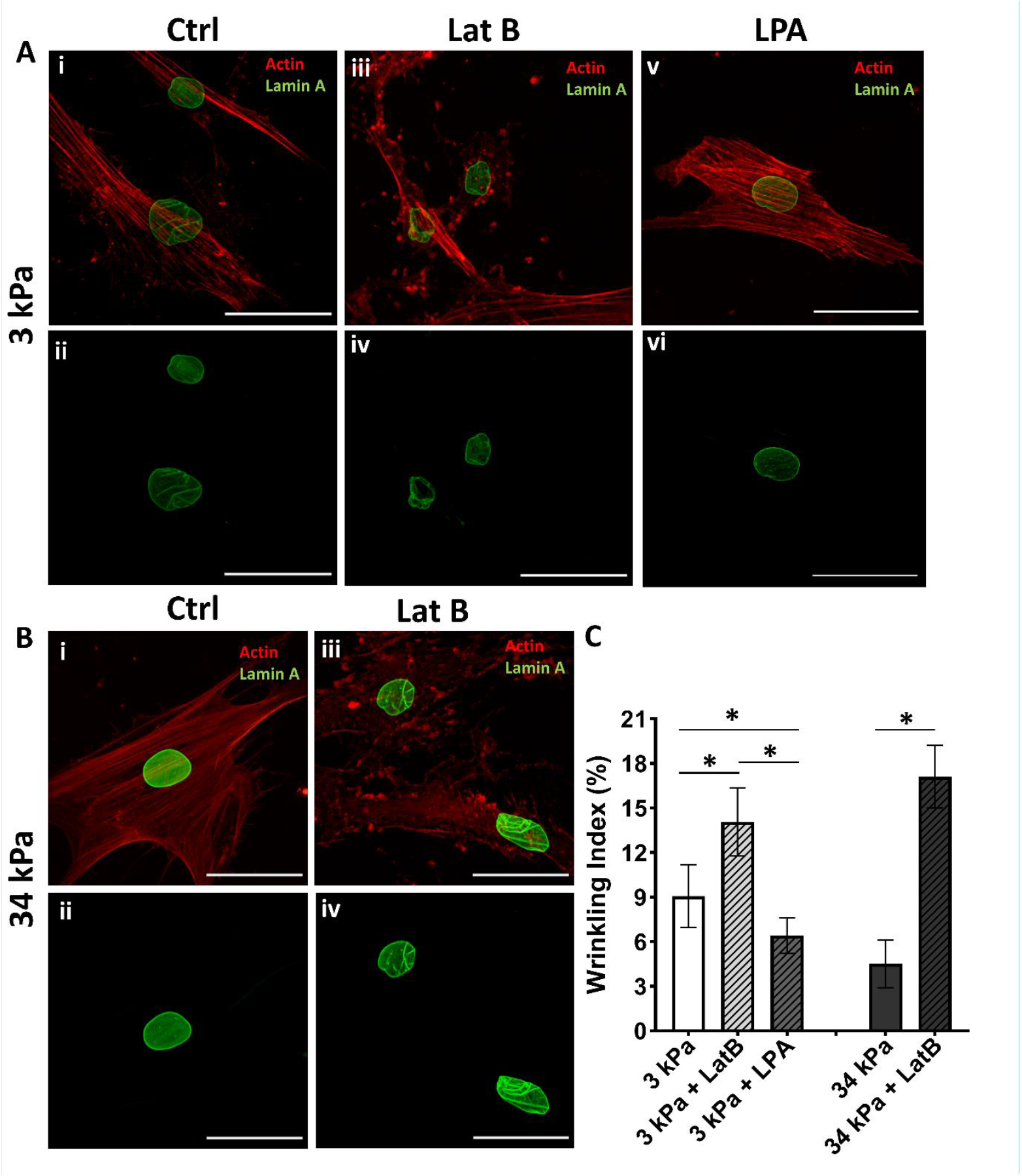
Effect of cellular contractility on nuclear wrinkling: Representative immunofluorescence images of actin (red) and lamin A (green) on (A) soft hydrogels (3 kPa) with control (i-ii), LatB (iii-iv), and LPA (v-vi), and (B) stiff hydrogels with control (i-ii) and LatB (iii-iv). (Graph C) shows quantification of wrinkling index on stiff and soft hydrogel with and without LatB and LPA. *p< 0.05, n>30 nuclei, scale bar: 50 µm.

**Fig. 8.**
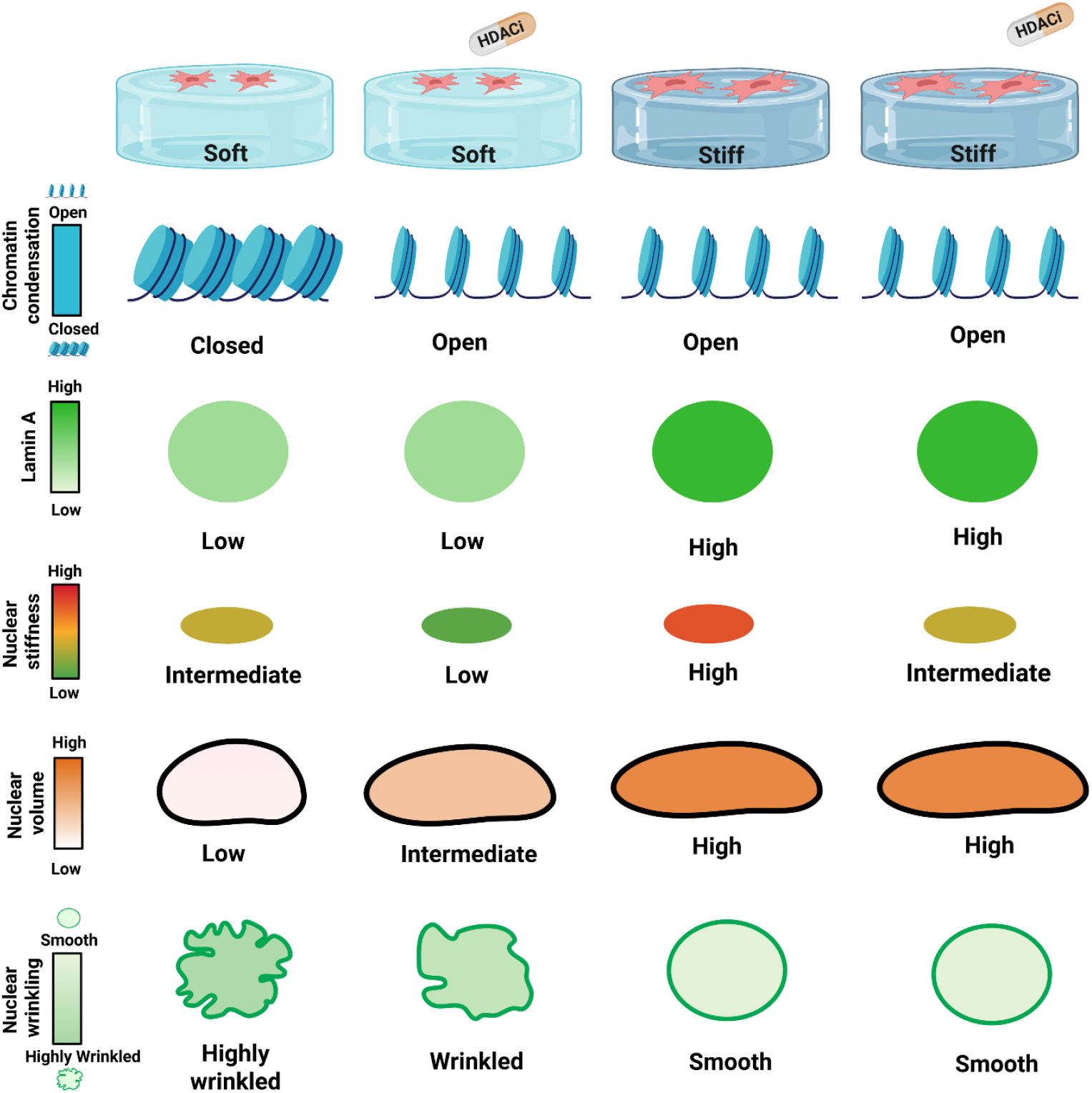
A summary showing effect of histone acetylation on nuclear morphology and architecture of hMSCs cultured on soft and stiff substrate: hMSCs in presence of HDACi mitigates the effect of soft substrate on nuclear morphological parameters including chromatin condensation and nuclear wrinkling while having minimal effect on hMSCs cultured on stiff substrate.

## Discussion

Over the past three decades, researchers have increasingly recognized the critical role of the mechanical microenvironment of tissues in regulating cellular behaviour, fate, and function^48,49^. Early investigations primarily focused on simple phenotypes such as cell spreading and migration. However, it is now well established that mechanosignaling affects a wide array of cellular processes, including differentiation, survival, secretion, and drug resistance^50^. Although the precise molecular mechanisms remain unclear, the nucleus has emerged as a central integrator of mechanical cues and biological outcomes.

In particular, the nucleus responds to mechanical stimuli through morphological alterations and changes in chromatin organization^26,51^. The significance of chromatin packing in mechanobiology has gained prominence, particularly in the context of disease pathology. In various pathological states, such as cancer and fibrosis, altered tissue stiffness is often accompanied by stiffness-dependent changes in cellular behaviors^52^. Nuclear mechanosensing and chromatin condensation have been identified as key mediators of these responses^53^. For example, in glioblastoma, stiffness-induced phenotypic changes have been shown to be regulated by nuclear mechanotransduction and associated chromatin remodeling.

Motivated by these insights, the present study investigated the combined influence of substrate rigidity and chromatin condensation on the nuclear morphometric features. Previously, our group demonstrated that the inhibition of histone deacetylases (HDACs) using valproic acid (VA) enhances cell spreading and modulates the mechanosensitive behaviour of human mesenchymal stem cells (hMSCs) on soft polyacrylamide (PAA) substrates^27^. Building on these findings, we examined how mechanical (substrate stiffness) and chemical (HDAC inhibition) cues together affect nuclear morphology and mechanics.

Our results show that VA-induced histone hyperacetylation on soft substrates leads to a significant increase in the projected nuclear area, volume, and surface area, with a corresponding reduction in the nuclear height (Fig.3). These morphological changes were not observed on stiff substrates, indicating that the nuclear response to HDAC inhibition is stiffness-dependent.

We further assessed nuclear stiffness, which is a key parameter in the cellular mechanobiology. Nuclear stiffness plays a critical role in maintaining the mechanical integrity and protecting the chromatin from mechanically induced damage. It also integrates mechanical or chemical cues from the microenvironment with chromatin organization, bridging the gap between physical forces and the nuclear architecture. Alterations in nuclear stiffness and chromatin organization are hallmarks of several diseases. In cancer, nuclei often exhibit reduced stiffness and increased deformability, facilitating migration through confined spaces during metastasis. These mechanical adaptations are frequently accompanied by global chromatin decompaction and epigenetic reprogramming processes. In this study, we investigated how substrate stiffness and HDACi, both known modulators of cellular mechanical and epigenetic states, affect nuclear stiffness. Interestingly, VA treatment resulted in decreased nuclear stiffness on both soft and stiff hydrogels. This finding contrasts with the effect of substrate stiffness, which is positively correlated with nuclear stiffness. Although both high substrate stiffness and VA treatment increased histone acetylation levels (Fig. 1), they had opposing effects on nuclear stiffness. These results suggest that substrate rigidity and HDAC inhibition regulate nuclear mechanics through distinct and possibly independent mechanisms.

Given that nuclear stiffness is influenced by both chromatin condensation and Lamin A expression, we examined the impact of VA and substrate stiffness on these factors. VA treatment significantly reduced chromatin condensation on both soft and stiff substrates (Fig.2), whereas Lamin A expression remained unchanged across all conditions(Fig.5). This indicates that the observed changes in nuclear stiffness are predominantly driven by chromatin decondensation, rather than alterations in nuclear lamina composition.

In addition to conventional nuclear morphometric parameters, we examined nuclear envelope wrinkling, a less frequently studied but pathophysiologically relevant feature. Cells cultured on soft hydrogels exhibited pronounced nuclear membrane wrinkling, which decreased with increasing substrate stiffness (Fig.6). Quantitative analysis revealed a stiffness-dependent decrease in the wrinkling indices. Notably, VA treatment significantly reduced nuclear wrinkling on soft substrates, whereas its effect was minimal on stiff substrates. These findings are consistent with recent reports, such as those by Mauck’s group^43^, suggesting that nuclear wrinkling may serve as a proxy for the mechanotransduction state of mesenchymal stem cells.

## Conclusion

In summary, this study demonstrates that chromatin deacetylation via HDAC inhibition substantially influences nuclear morphology and mechanics, particularly under low-stiffness conditions. These changes appear to be largely independent of lamin A and are mediated through alterations in chromatin organization. These findings open promising avenues for future research aimed at screening small-molecule modulators, including HDAC inhibitors (HDACi), histone acetyltransferase inhibitors (HATi), and histone methyltransferase inhibitors (HMTi), to better understand their impact on nuclear morphology and function. An important question arising from this study is whether nuclear deformations resulting from matrix dysregulation in diseases can be reversed by such interventions. We hypothesize that preserving nuclear morphology through chromatin-level regulation may offer a potential intervention strategy for various diseases and aging.

## Author Information

### Authors

**Rohit Joshi**- Department of chemical Engineering, Indian institute of technology Bombay, Maharashtra 400076.

**Darshan Shah-** Department of chemical Engineering, Indian institute of technology Bombay, Maharashtra 400076.

**Venkatesh V Kareenhalli-** Department of chemical Engineering, Indian institute of technology Bombay, Maharashtra 400076.

## Author Contribution

A.M. and R.J. conceptualized and designed the research; A.M. supervised the project. R.J., D.S. performed the experiments. R.J., D.S. analysed the data. R.J. prepared the figures. R.J. and A.M. interpreted the results. R.J., A.M. drafted the manuscript. A.M., V.K. provided the critical inputs on the manuscript.

## Conflict of interest

The authors declare no conflict of interest.

## Data Availability

All relevant data and associated protocols are provided within the article and *SI Appendix*.

## Acknowledgement

RJ thanks DBT for the fellowship (DBT/2016/IIT-B/560). We thank SERB-CRG research funding with grant no. CRG/2022/005882 and WRCB-IITB: DO/2018-WRCB002-021. We thank IIT Bombay for infrastructural support, laser scanning confocal microscopy central facility and atomic force microscopy central facility.

## Notes

### Competing Interest Statement

The authors have declared no competing interest.

