## Supplementary information for "Histone acetylation alters nuclear morphology and architecture of human mesenchymal stem cells in rigidity dependent manner"

Table S1

| Acrylamide from 40% stock solution (ml) | Bis-acrylamide from 2% Stock solution (ml) | Water (ml) | E ± St. Dev. (kPa) |
|---|---|---|---|
| 1 | 1.125 | 7.875 | 3.13 ± 0.42 |
| 2.5 | 1.5 | 6 | 34.88 |

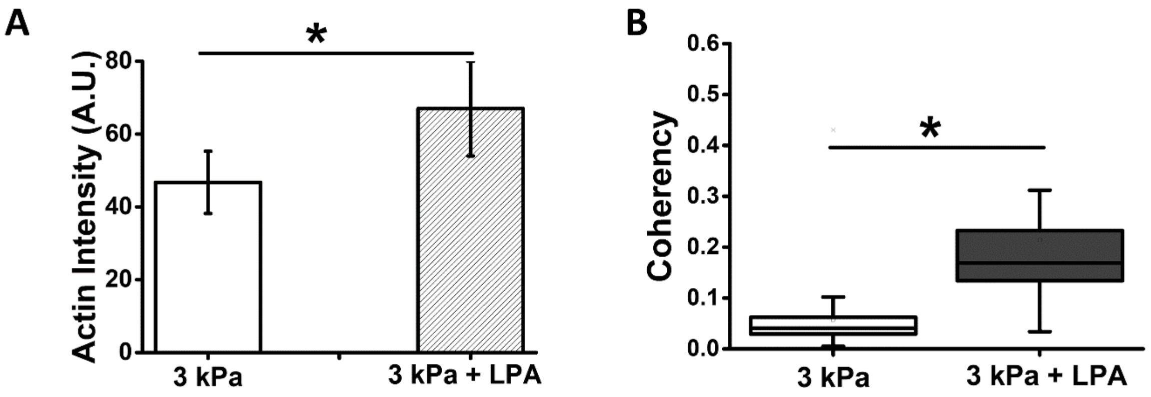


**Fig. S1.** Quantification of Actin intensity (A) and actin coherency (B) on soft hydrogel with and without LPA. *p<0.05, n>30.
